# Unsupervised phenotypic analysis of cellular images with multi-scale convolutional neural networks

**DOI:** 10.1101/361410

**Authors:** William J. Godinez, Imtiaz Hossain, Xian Zhang

**Affiliations:** Novartis Institutes for BioMedical Research, Basel, Switzerland.; Current address: Novartis Institutes for BioMedical Research, Emeryville, CA, USA

## Abstract

Large-scale cellular imaging and phenotyping is a widely adopted strategy for understanding biological systems and chemical perturbations. Quantitative analysis of cellular images for identifying phenotypic changes is a key challenge within this strategy, and has recently seen promising progress with approaches based on deep neural networks. However, studies so far require either pre-segmented images as input or manual phenotype annotations for training, or both. To address these limitations, we have developed an unsupervised approach that exploits the inherent groupings within cellular imaging datasets to define surrogate classes that are used to train a multi-scale convolutional neural network. The trained network takes as input full-resolution microscopy images, and, without the need for segmentation, yields as output feature vectors that support phenotypic profiling. Benchmarked on two diverse benchmark datasets, the proposed approach yields accurate phenotypic predictions as well as compound potency estimates comparable to the state-of-the-art. More importantly, we show that the approach identifies novel cellular phenotypes not included in the manual annotation nor detected by previous studies.

**Author summary:** Cellular microscopy images provide detailed information about how cells respond to genetic or chemical treatments, and have been widely and successfully used in basic research and drug discovery. The recent breakthrough of deep learning methods for natural imaging recognition tasks has triggered the development and application of deep learning methods to cellular images to understand how cells change upon perturbation. Although successful, deep learning studies so far either can only take images of individual cells as input or require human experts to label a large amount of images. In this paper, we present an unsupervised deep learning approach that, without any human annotation, analyzes directly full-resolution microscopy images displaying typically hundreds of cells. We apply the approach to two benchmark datasets, and show that the approach identifies novel visual phenotypes not detected by previous studies.

## Introduction

Image-based high-throughput cellular assays allow meticulous monitoring of chemical or genetic perturbations of cellular systems at large scale(1–4). Quantitative analysis of the collections of image data generated by these assays is pivotal for an objective assessment of the phenotypic diversity observed within the data. Conventional workflows developed for image analysis involve a series of disjoint data-processing tasks, such as detection of cellular objects, numerical characterization of these objects via feature engineering, as well as classification of cellular objects based on their features into different phenotypes(5,6). Many of these steps have been addressed with the *deep learning* methodology(7,8), which has previously yielded state-of-the-art results for such computer vision tasks(9–13). Approaches(14–16) based on deep learning for analyzing high-content cellular images follow primarily a *supervised learning* paradigm, whereby images annotated with phenotypic labels are used to train a deep neural network model that maps images to one of the labels. The predictions of supervised approaches are therefore constrained to the set of phenotypes defined during training, and therefore do not naturally support the identification of additional phenotypes. The acquisition of these phenotypic labels through manual annotation of the image data is also time-consuming (e.g., requiring crowdsourcing efforts(17)), and error-prone(18). The applicability of supervised approaches is thus contingent upon the availability and quality of the manual annotation.

Strategies to escape the limitations imposed by the a priori definition and acquisition of phenotypic labels include *transfer learning* as well as *unsupervised learning*. In the former, a neural network classification model trained in a supervised manner on a non-cellular image dataset is applied to a cellular image dataset(19). Since the categories defined in the *source* non-cellular dataset do not match those of the *target* cellular dataset, the aim of this strategy is to map cellular images to a continuous coordinate system, i.e., a *feature space*, by treating the activation of the hidden layers of the pre-trained deep model as a feature vector. While this strategy has been shown to work well for extracting biologically informative features(19), there are no guarantees that models trained on non-cellular data generalize well to arbitrary cellular image data. Technical issues such as different channel encodings (e.g., RGB channels in non-cellular images compared with an arbitrary number of fluorescence channels in cellular images) and noise models (e.g., additive Gaussian noise models in non-cellular images(20) compared with Mixed-Poisson-Gaussian statistics(21) in fluorescence images) also hinder the applicability of approaches based on transfer learning.

Approaches following an unsupervised learning paradigm are, in contrast, typically optimized on the specific cellular dataset of interest. The aim of unsupervised learning is to map images to a feature space where biologically relevant patterns within the dataset might emerge. While in the supervised learning paradigm deep models are designed to predict an *extrinsic* characteristic or attribute of the data, e.g., the phenotypic label manually assigned to the images, in the unsupervised learning paradigm deep models are designed to predict an *intrinsic* characteristic of the data. The most inherent property of each image is the pixel data itself. The training process of both *autoencoder networks*(22) as well as *generative adversarial networks* (23)(GANs) therefore typically involves the optimization of an image synthesis function aiming to reconstruct an image’s raw pixel data from a low dimensional representation of the input image. This type of approaches has been able to map single-cell images with small dimensions (e.g., 40 × 40 pixels) to a low-dimensional space (e.g., 64-D) where aberrant morphologies during cell division as induced by siRNAs may be identified(24). Because of the high spatial and phenotypical variability found in multi-cellular images with larger dimensions (e.g., 1280 × 1024 pixels over three fluorescent channels, i.e., a 3932160-D space), the compression and reconstruction of high-content images via a neural network is currently a computationally prohibitive task.

Here we present an unsupervised approach based on the *exemplar convolutional neural network(25)* (Exemplar-CNN) training methodology that optimizes a network model to discriminate among *surrogate classes*, which, in our case, are automatically defined through the intrinsic groupings of images (e.g., images belonging to the same treatment) typically found in high-content imaging studies. The proposed approach uses exclusively dataset-specific multi-cellular images, and requires no phenotypic annotations or optimization of computationally-expensive image reconstruction functions. Using this unsupervised strategy, we train our multi-scale convolutional neural network architecture (M-CNN(16)) on the multi-cellular images of the KiMorph(26) and BBBC021(27,28) datasets, which involve genetic and chemical perturbations of cellular systems at scale, respectively. Our approach, without any user-provided phenotypic labels and without any object segmentation, is able to map images to a feature space that enables prediction of phenotypes that match well with held-out labels. In addition, we show that the approach identifies novel phenotypes in the benchmark datasets not detected by previous studies.

## Results

### Training and validating a deep neural network with the Exemplar-CNN methodology

To train a neural network model without any user-provided phenotypic annotation, we used the Exemplar-CNN(25) training methodology (see **Methods** for details). When applied to high-content cellular images, the trained network model maps in one step an image to a continuous feature space representing the phenotypic homogeneity and variability observed in the data (see **Fig. 1** for a schematic overview of the approach). We validated this unsupervised strategy on the Kimorph and BBBC021 datasets, which include images of cells subjected to siRNA and compound treatments, respectively. On each dataset, we trained a multi-scale convolutional neural network (M-CNN(16)) model with the Exemplar-CNN training methodology. To define the surrogate classes required by this methodology, we hypothesized that images belonging to the same well (KiMorph) or compound treatment (BBBC021) defined a single surrogate class. No phenotypic categories and annotations were therefore needed for training. Note that different surrogate classes might belong to the same phenotypic category, but this information is not known to the network. We trained the neural network exclusively with the pixel data of annotated images; the annotations were removed during training, and only used subsequently to validate the performance of the approach.

**Fig. 1.**
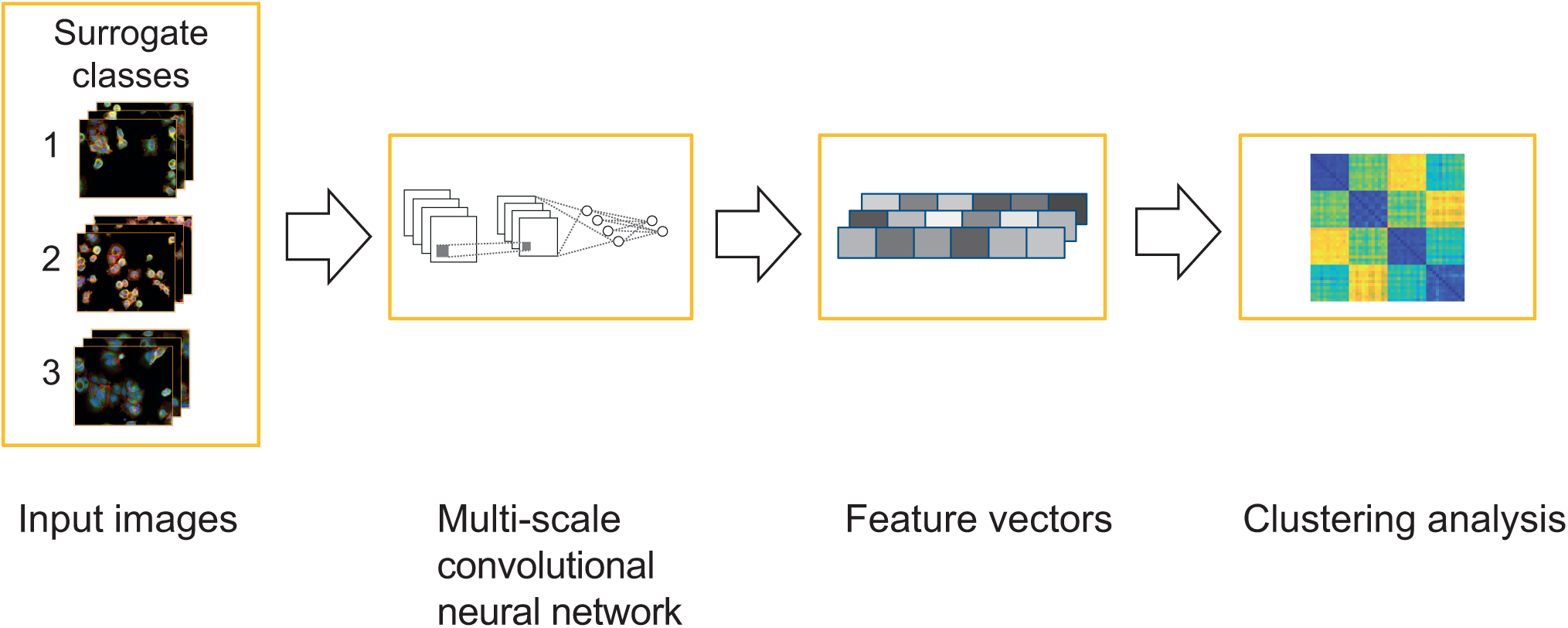
Schematic overview of the Exemplar-CNN approach. Images are grouped into surrogate classes based on intrinsic information (such as treatment information) instead of external annotation. Taking the full-resolution images of the surrogate classes as input, an M-CNN model is trained with the objective of separating the different surrogate classes. Once trained, as input images are fed to the network, the neural activation values are extracted as feature vectors, thus mapping the input images to a low-dimensional feature space. Finally, distance calculation and clustering analysis enable the identification of novel phenotypes.

Once the dataset-specific network models were trained, we validated the performance of the approach in two steps. First, we built a nearest-neighbor classifier based on the feature vectors computed by the trained M-CNN model to predict the phenotype of annotated images. We used the classification accuracy of the nearest neighbor classifier to evaluate whether the feature vectors computed by the network encoded relevant phenotypic information. Second, we applied the trained M-CNN model to the dataset’s entire image collection, including images with no annotation and not used during training, thereby obtaining one feature vector for each image in the entire collection. We performed hierarchical clustering analysis on the feature vectors, and, through visual inspection of images in selected clusters, identified novel phenotypes. The next two sections describe in detail the results for each dataset.

### KiMorph analysis and results

The Kimorph dataset comes from an RNAi screen where HeLa cells were reverse transfected with siRNAs targeting ca. 800 kinases in duplicate. siRNA-mediated perturbations of the UBC, CLSPN and TRAPPC3 genes were used as positive controls while the Renilla luciferase (*Rluc*) siRNA treatment was used as a neutral control. After transfection, cells were fixed and labeled for DNA, F-actin, and B-tubulin, and imaged through an automated microscope (experimental details can be found in the original publication(26)). Each of the four control siRNA treatments (UBC, CLSPN, TRAPPC3, and Rluc) was spotted across 12 wells in duplicate. We declared all fields-of-view (FOVs) coming from each well to define a single surrogate class, which amounted to 48 surrogate classes (i.e., one class per replicate well). We then trained an M-CNN model to maximize separation among these classes (see **Methods** for training details). Note that some surrogate classes belong to the same control siRNA treatment but this information is not known to the network. Previous studies(26,29) have shown that each of the four control siRNA treatments induces a consistent phenotype across wells. If the network learned to identify invariant and discriminative features reflecting the control phenotypes, we hypothesized that these would remain relatively similar within surrogate classes belonging to the same control while varying more strongly across surrogate classes belonging to different controls.

After training, we used the M-CNN model and PCA to extract a 94-dimensional feature vector for each original FOV image. We aggregated the feature vectors at the well level, and calculated cosine distance values between each pair of wells (see **Methods** for details). The resulting distance matrix, with rows ordered by control groups, is shown in **Fig. 2a**. We observe square blocks (submatrices) along the diagonal of the matrix, which reflect the low distance values (i.e., high similarity) within wells from the same control siRNA treatment as entailed by the feature vectors computed by the network. To quantitatively verify the invariant and discriminative properties of the feature vectors within and across the four different phenotypes, we tested whether we could predict the phenotype of each well based on the phenotype of the well’s nearest neighbor in feature space. Over 50 repetitions of a random hold-out cross-validation strategy, this nearest-neighbor classification approach identifying the four control phenotypes yielded 100% classification accuracy (**Supplementary Table 1**). The results show that the feature vectors computed by the M-CNN model, which was trained without any phenotypic annotation, remain relatively similar within the same phenotype yet vary across different ones, thus encoding phenotypic information that enables the identification of distinct phenotypes.

**Fig. 2.**
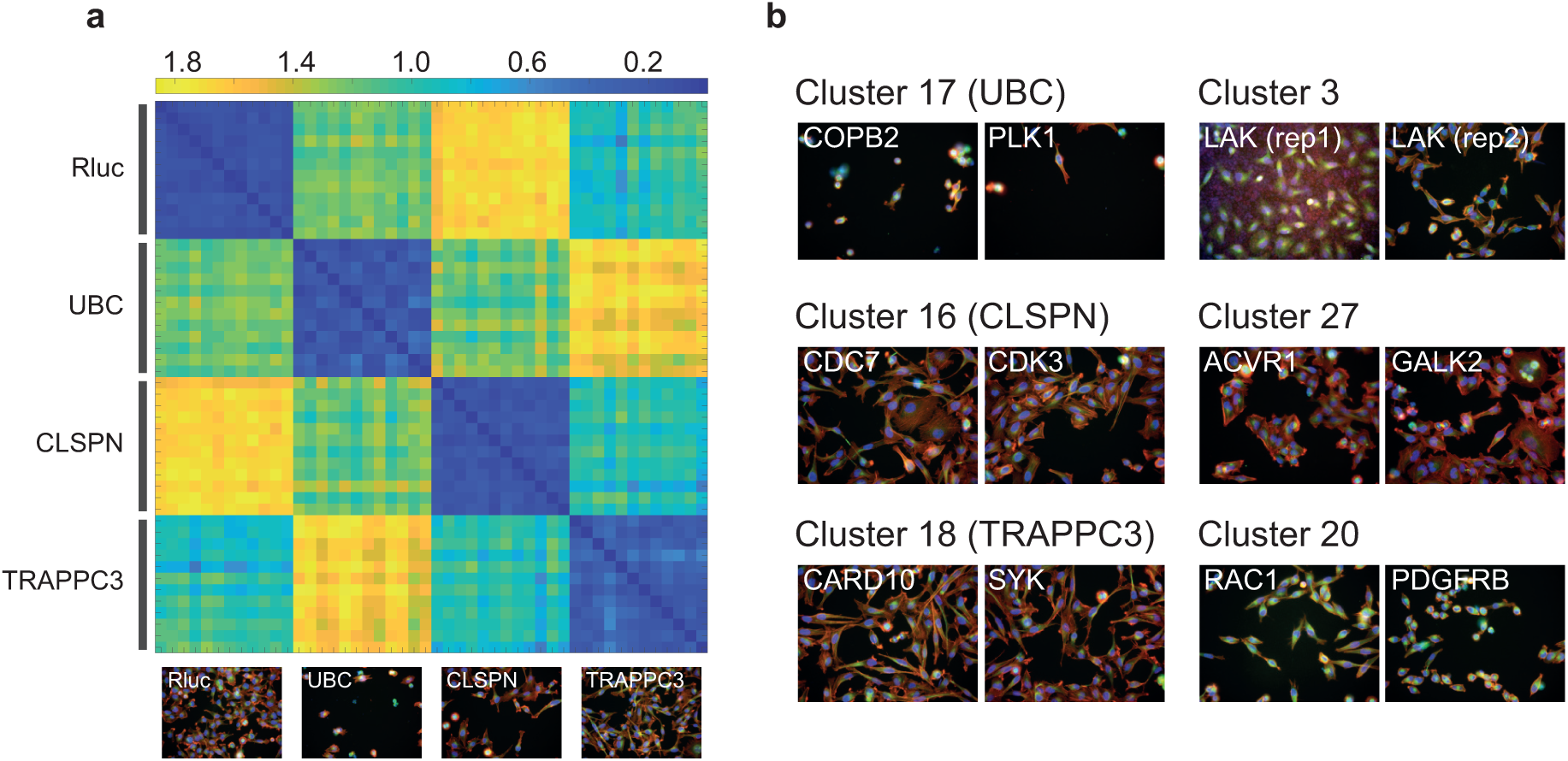
Exemplar-CNN training and clustering analysis results for the KiMorph dataset. (**a**) Cosine distance matrix between pairs of wells of control siRNA treatments. Rows are ordered by control groups. Blue indicates a small distance value while yellow corresponds to a large distance value. (**b**) Sample images from clusters including the control siRNA treatments (left column) as well as from clusters including phenotypes distinct from the control treatments (right column).

We next tested whether the M-CNN model trained with four control siRNA treatments could generalize to images and phenotypes not used during training. We therefore fed each image of the entire KiMorph dataset through the trained M-CNN model and PCA, and obtained a 94-D feature vector per image. Feature vectors of images belonging to the same siRNA treatment were aggregated onto a single vector (see **Methods** for details). We calculated pairwise cosine distances among all vectors corresponding to all 781 siRNA perturbations (see **Supplementary Table 2**), performed hierarchical clustering, and grouped all siRNA perturbations onto 27 clusters (see **Supplementary Table 3**). On each cluster, we carried out a Gene Ontology (GO) enrichment analysis for biological processes through the topGO R package(30), with all kinases in the library taken as the background set. After Benjamini-Hochberg correction for multiple testing, we found that 22 clusters out of the 24 clusters that included more than one gene were enriched with two or more GO terms. This indicates that the feature vectors computed by the network supported the identification of shared biological functions of groups of genes (see **Supplementary Table 4**). To validate the results, we first inspected clusters 17, 16, and 18, which included the three positive siRNA controls (viz. UBC, CLSPN, and TRAPPC3), respectively (**Fig. 2b**). The UBC cluster, which the enrichment analysis associates with DNA damage and integrity checkpoints, includes essential genes such as COPB2 and PLK1 that, when knocked down, cause a lethal phenotype akin to that of the UBC treatment. Likewise, the CLSPN cluster comprises genes such as CDC7 and CDK3, which are associated with cell cycle control, and whose knockdowns induce an enlarged cell phenotype resembling that of the CLSPN treatment. In the TRAPPC3 cluster, which is enriched with the ‘integrin-mediated signaling pathway’ GO term, we typically observe an elongated cell phenotype that is likewise triggered by the CARD10 and SYK knockdowns. Overall, the clustering results and the recapitulation of known biological functions of groups of genes suggest the feature vectors learned by the network capture phenotypic information.

The main advantage of an unsupervised approach is its ability to discover novel phenotypes. To verify this premise, we identified clusters that were relatively distant (in terms of the cosine distance values) from the four siRNA controls (see **Methods** for details). One of these distant clusters (viz. cluster 3) only includes images from the LAK perturbation. Visual inspection of the images reveals an experimental artifact in images from replicate 1 (**Fig. 2b** top right; compare to images from replicate 2). The artifact is not visible in images of any other treatment, and so the approach correctly grouped images with this artifact onto a separate cluster. Images from cluster 27, which includes genes such as ACVR1 and GALK2, display a phenotype of enlarged nuclei and cells with a strong actin signal (**Fig. 2b** middle right) that do not resemble any of the control phenotypes. Likewise, in cluster 20, which includes genes such as RAC1 and PDGFRB, and shows enrichment for the ‘positive regulation of Rho protein signal transduction’ GO term, we observe a reduced cell count and cell size in the images. These results suggest that the proposed unsupervised strategy supports the identification of novel imaging phenotypes.

### BBBC021 analysis and results

In the BBBC021 dataset, MCF-7 breast cancer cells were treated with 113 compounds at eight concentrations in triplicate, before being fixed and labeled for DNA, F-actin, and B-tubulin. Images were captured from each channel with four fields per well(28). A subset of 103 compound-concentration pairs (hereafter defined as treatments) covering 38 compounds was previously inspected and annotated for one of twelve mechanisms-of-action (MoAs)(31). We declared all images coming from each treatment to belong to the same surrogate class, which resulted in 103 surrogate classes. The M-CNN model was trained to discriminate among all 103 surrogate classes (see **Methods** for training details). Note that the network is not aware that certain surrogate classes (i.e., treatments) belong to the same compound or the same MoA. If the network learned MoA-relevant features, we posited that these would remain relatively invariant within each MoA yet vary across different MoAs.

Once trained, we used the M-CNN model and PCA to extract an 8-D vector for each input image used during training. We aggregated the feature vectors of images belonging to the same treatment onto a single feature vector and computed cosine distances between all pairs of treatments (**Fig. 3a)**. Each row of the matrix corresponds to one treatment. Treatments are ordered by MoA, and then by compound and concentration. Overall we observe sub-matrices (squares) along the diagonal indicating that the network learned features that remain relatively invariant within each MoA. To quantitatively verify the homogeneity and variation of the learned features within and across phenotypes, respectively, we tested whether the MoA of a treatment could be identified based on the MoA label of the treatment’s nearest neighbor in feature space. Here we adopted the same leave-one-compound-out cross-validation strategy used in previous benchmarking studies(31) that prevents matching treatments from the same compound (see **Methods** for details). With this nearest-neighbor classification strategy, we achieve a median accuracy over all classes of 88%. The confusion matrix is shown in **Supplementary Table 5**. For certain MoAs (e.g., aurora kinase inhibitors, cholesterol-lowering, protein degradation, and protein synthesis), the approach achieved 100% classification accuracy. For other MoAs, the accuracy ranged from 75% to 89%. The performance of the approach is comparable to our previous supervised approach(16), which is explicitly optimized to distinguish these twelve MoAs, as well as to other non-supervised approaches(19,31). Overall the results show that, while the MoA categories are unknown to the network, it manages to learn features which remained relatively invariant within each MoA yet varied across MoAs.

**Fig. 3.**
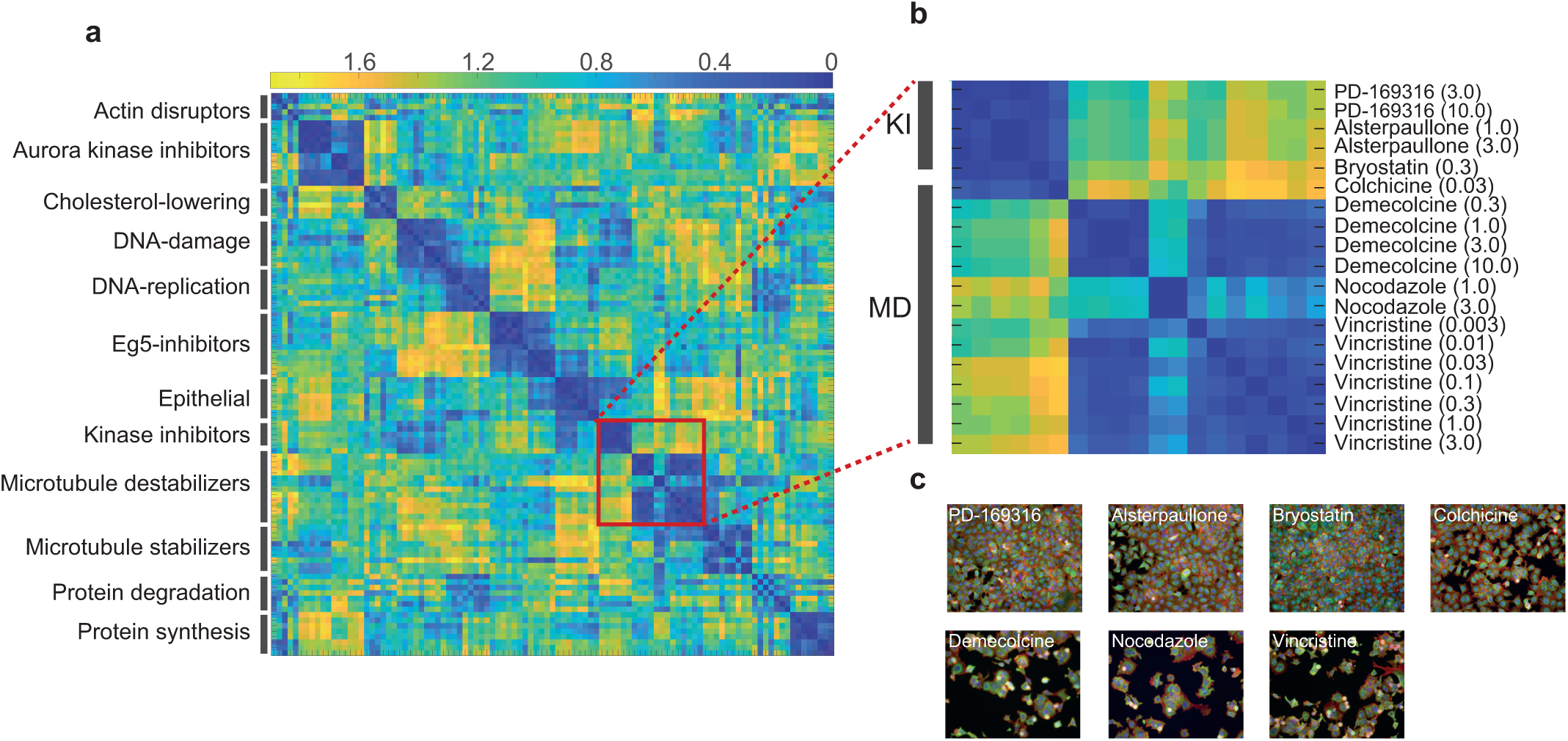
Exemplar-CNN training results for the BBBC021 annotated subset. (**a**) Cosine distance matrix between pairs of treatments. Labels on the left are the MoA annotations. Rows are ordered by MoA annotation, compound, and concentration. Blue indicates a small distance value while yellow corresponds to a large distance value. (**b**) Zoomed-in view of the red box in (a), KI for kinase inhibitors and MD for microtubule destabilizers. Labels on the right are compounds and concentrations in μM. (**c**) Sample images corresponding to the treatments in (b). Colchicine, which is annotated as a microtubule destabilizer (MD), visually looks more similar to treatments annotated as kinase inhibitors (KI).

While the approach is able to recapitulate known information about the annotated data, we tested further its ability to reveal phenotypic information beyond the annotation. To this aim, we conducted a closer examination of the distance matrix. While treatments annotated with the same MoA are mapped by the network to nearby positions in feature space, and are therefore distinguishable via a nearest-neighbor classifier, the sub-matrices along the diagonal of the distance matrix in **Fig. 3a** reveal a certain heterogeneity within individual MoAs. For example, for the aurora kinase inhibitors (Aur) MoA, the corresponding sub-matrix reveals three groups corresponding to the three compounds annotated with this MoA (viz. AZ-A, AZ258 and AZ841), which suggests that the compounds caused slightly different sub-phenotypes. A similar observation can be made for the actin disruptors (Act), protein degradation (PD), and protein synthesis (PS) MoAs. The microtubule destabilizers (MD) MoA is comprised by four compounds (14 treatments, sub-matrix highlighted in red in **Fig. 3a**. and zoomed in view in **Fig. 3b**). Three of the four compounds, Demecolcine, Nocodazole, and Vincristine, are relatively similar to each other as well as distant to the kinase inhibitor (KI) group, although Nocodazole shows a sub-phenotype different from Demecolcine and Vincristine. The fourth compound, Colchicine at 0.03μM, which is annotated as MD, instead seems to be closer to the kinase inhibitors (KI) treatments than to the other MD treatments, and is accordingly predicted as KI by the nearest-neighbor classification scheme. Visual examination of the corresponding images also confirms Colcichine’s similarity to the KI treatments (**Fig. 3c**). Although only one concentration (0.03μM) of Colcichine is included in the annotation subset, there are seven concentrations (0.001 – 3.0 μM) in the entire BBBC021 dataset. Only at 3.0μM, Colchicine causes phenotypes similar to other microtubule destabilizers (**Supplementary Fig. 2**). The proposed unsupervised approach is thus able to detect phenotype information in the data beyond the manual annotation, which is not feasible with a supervised method.

Next, we set out to verify the applicability of the approach to the entire BBBC021 dataset. Here we first trained an M-CNN model using all images from the 103 annotated treatments (amounting to 103 surrogate classes) plus images from the neutral control (DMSO), which were grouped into an additional surrogate class. Once trained, we applied the model to all 13200 images included in the dataset. Using PCA, we obtained a 77-D feature vector per image. Vectors of images belonging to the same treatment replicate (well) were aggregated onto a single vector. We then determined the *similarity* of each replicate vector to each of the 12 MoAs and DMSO based on the cosine distances of each vector to all replicate vectors belonging to the 103 MoA-annotated treatments as well as to all DMSO wells (see **Methods** for details). The 13 similarity values of each treatment replicate are shown in **Supplementary Table 6**.

In our previous supervised analysis of the BBBC021 dataset, four compounds were selected as representative concentration-response curves (16). In the current study, we selected the same four compounds and plotted the similarity values to each MoA and DMSO as a function of the concentration (**Supplementary Fig. 1**). In the previous supervised approach, the y-axis was the classification probability and for each compound there was only one or two dominant MoAs across concentrations. In the current unsupervised approach, the y-axis is the similarity to each MoA which ranges from 0 to 2, and the gaps between curves are much less pronounced. To compare the overall trend, we simplified the plot by only showing MoAs shown as dominant in the previous supervised approach (**Fig. 4a**). Data points highlighted by dashed circles correspond to concentrations annotated with the curve’s MoA (and therefore achieving maximum similarity). For Floxuridine, consistent with the supervised approach, the DNA replication (DR) MoA is the top MoA prediction for all concentrations. For Nocodazole, the curve shows a similar trend to the supervised approach, with DMSO as the top MoA in low concentrations (0.001-0.01μM) and microtubule destabilizers (MD) as the top MoA in high concentrations (0.1-4.0μM). For Alsterpaullone, in the current unsupervised analysis, kinase inhibitor (KI) and DMSO are on the same level until the higher concentrations. DNA-damage (DD) increases at the last concentration but does not pass the level of KI. In the previous supervised analysis the differences among the MoAs were much more obvious although with larger error bars. Finally, for Hydroxyurea, for which none of the concentrations was included in the training data, the trend of the curve is consistent with the supervised approach, where DMSO decreases over concentration while DNA-damage (DD) increases and takes over at the two highest concentrations.

**Fig. 4.**
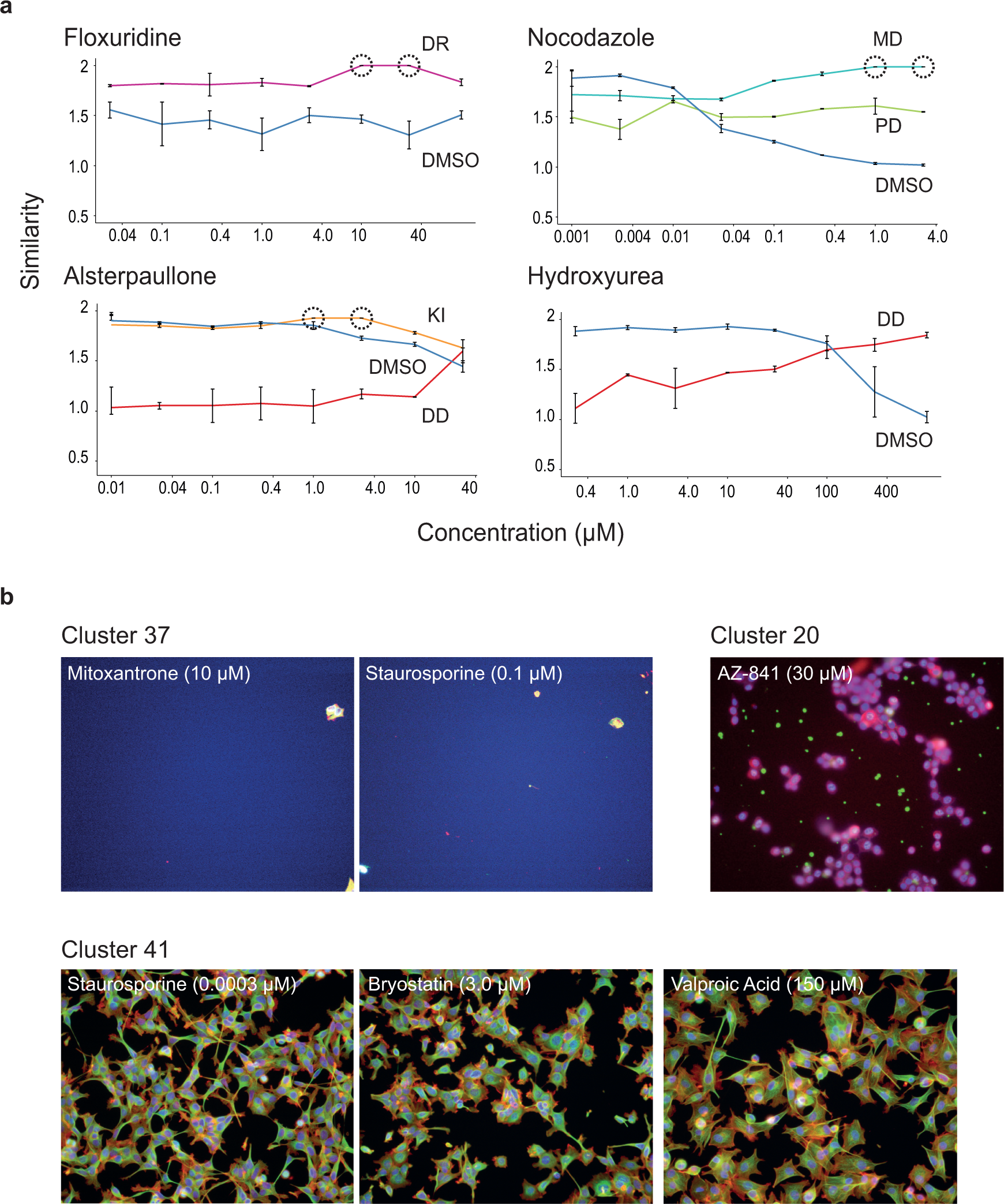
Example concentration-response curves and clustering analysis for the BBBC021 dataset. (**a**) Similarity-vs-concentration plots for four compounds. The similarity (y-axis) to selected MoAs and DMSO over concentration (x-axis) computed using the features vectors yielded by the proposed approach is shown. The dots and error bars represent the median and MAD over the experimental replicates (*n* = 2 for Alsterpaullone and *n* = 3 for the other three compounds). Data points marked by dashed circles are annotated with the curve’s MoA and therefore achieve maximum similarity. (**b**) Sample images of clusters including distinct phenotypes not related to the annotated phenotypes.

Finally, we tested the ability of the approach to detect novel phenotypes. To this end, we further aggregated the replicate vectors belonging to the same treatment onto a single vector. Likewise, DMSO vectors stemming from the same plate were aggregated onto a single vector. We calculated pairwise cosine distances among all treatments, including DMSO, and applied a hierarchical clustering procedure that yielded 79 clusters (see **Methods** as well as **Supplementary Table 7** for the complete distance matrix). We inspected visually clusters that included exclusively compound-concentration treatments without any MoA annotation (see **Fig. 4b**). For example, in cluster 37, we found images from Mitoxantrone at 10μM and Staurosporine at 0.1μM and 0.3μM that induced a strong toxic phenotype. In cluster 20, which included images from AZ-841 at 30μM only, we found images that displayed an unusual purple phenotype that could hint at a tubulin toxin/disruptor MoA for this treatment. Finally, in cluster 41, we found images from Staurosporine at 0.0003μM, Bryostatin at 3.0μM, as well as Valproic Acid at 150μM where groups of elongated cells with thin protrusions forming a networked pattern were visible. The results underscore the ability of the proposed unsupervised approach to identify novel phenotypes not previously known and not included during training.

## Discussion

Deep learning has been successfully pioneered in the field of image-based high-throughput screening(14–16,19,24). The majority of approaches based on deep neural networks adopt a supervised learning paradigm that requires manual definition and acquisition of phenotypic labels. As such, supervised approaches do not support naturally the discovery of new phenotypes. In this work, instead of relying on predefined phenotypic labels, we developed an unsupervised learning approach that exploits the inherent variation across treatments typically found in imaging-based studies to learn phenotypically relevant features that enable the discovery of novel phenotypes.

The proposed approach obviates the need for manually specified phenotypic categories by defining automatically surrogate categories through the inherent grouping of images (e.g., images belonging to the same well) found in the experimental design of high-content studies. The fact that multiple surrogate categories may belong to a (known) phenotypic class remains explicitly held-out to the neural network model throughout. Our results on two benchmark datasets demonstrate that the feature vectors extracted from the images through the trained models support the recognition of known phenotypes included within the surrogate categories. By testing the models on images outside of the surrogate categories, we also showed that the models generalize to phenotypes beyond those used during training. With a straightforward clustering analysis of the feature vectors, we managed to pinpoint novel phenotypes, which is one of the main goals of image-based high-content screening studies, where genetic or chemical perturbations may potentially induce a range of unexpected phenotypes.

Certainly, one could identify novel phenotypes with conventional image analysis approaches, which typically require segmentation and manual feature engineering(26,28,32,33). It is however encouraging to see that the proposed unsupervised approach, which requires no segmentation, no manual feature engineering, and no phenotypic categories and annotations, also supports the identification of novel phenotypes in a more automated fashion. The proposed approach does not provide single-cell readouts, and therefore does not replace single-cell analyses(34,35).

With the proposed unsupervised learning strategy, the inferred network models depend on the phenotypic data included within the surrogate classes. In our study, we restricted the surrogate classes to images that had a phenotypic annotation. This strategy facilitated the validation of the approach, as it allowed testing whether the approach supported the recovery of known phenotypic classes. Additional work is however needed to decide which images and phenotypes should be included within the surrogate classes. One possibility would be to adapt an active learning approach, where surrogate classes would be iteratively added based on a certain performance criterion.

## Methods

### Exemplar Convolutional Neural Networks

We use the Exemplar-CNN optimization strategy(25) to train a convolutional neural network without relying on any phenotypic label annotation. In contrast to a typical supervised learning approach, where the neural network is trained to discriminate among a set of predefined phenotypic classes, the proposed approach is trained to discriminate among a set of *surrogate classes*. The main idea underlying the Exemplar-CNN methodology is to learn image features that are both *invariant* within each surrogate class as well as *discriminative* across surrogate classes. In the original strategy, each *exemplar* (i.e., a region-of-interest within an image) and transformed versions thereof (obtained through extreme data augmentation schemes) defined a single surrogate class. This strategy was shown to work well with a large number of surrogate classes (e.g., up to 4000). However, when the number of (exemplar) images is very large and the images look very similar, discrimination among the surrogate classes becomes more challenging. A prior grouping (e.g., through clustering) of similar images was suggested as an approach to reduce the number of classes as well as to group very similar images into a single surrogate class.

In our case, instead of taking each image and its variations as a single surrogate class, we take advantage of the intrinsic grouping of images provided by the experimental design of each study to define the surrogate classes. For example, for each well, multiple fields-of-view (FOVs) are typically acquired. We may therefore define all FOVs from a single well to define a single surrogate class. Similarly, each treatment combination (e.g., a compound at a specific concentration) is typically replicated. Images from these replicates may be therefore declared as a single surrogate class. The definition of surrogate classes depends on the experimental details in each study. After defining *N_s_* surrogate classes in such a way, we associate a numerical label *y*_surrogate_ with each surrogate class and its images.

We use a multi-scale convolutional neural network (M-CNN) architecture to solve the task of surrogate class discrimination. The last two layers of our M-CNN architecture include a fully connected layer with 128 hidden units, as well as a soft-max output layer, which yields a vector **ρ** with elements *ρ_k_* that encode a probability score for each of the *N_s_* surrogate classes to be identified (all architectural details are provided in **Supplementary Table 8**). Using *N_t_* images associated with surrogate classes and their numerical labels, we optimize the parameters of the M-CNN by minimizing the following error function:

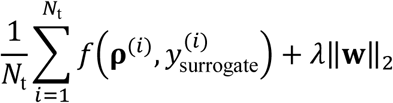

where *f(*⋅,⋅) is the cross-entropy error function evaluating the agreement between the network’s soft-max output **ρ**^(i)^ and the surrogate (true) label 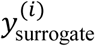 for the *i*-*th* training example, ∥⋅∥_2_ is the L2 norm, **w** is a vector including all weights of the network, and *λ* is a coefficient that regulates the influence of the magnitude of the weight vector on the error function. We use the stochastic gradient descent (SGD) algorithm via backpropagation and drop-out to approximate a solution.

### Learning details

Generally, we used the same strategy and parameter values that we used previously to train the M-CNN architecture in a supervised way(16). In this study, we however increased the number of training epochs to 27. The step size over which the learning rate is held constant was also increased to 9 epochs. One epoch is equal to the number of iterations needed to evaluate all images in the training dataset. We additionally used the dropout technique(36) on the penultimate layer of the M-CNN architecture to encourage a better exploration of the available activation space.

### Feature extraction, projection, and aggregation

Once trained, the application of the M-CNN model to any input image yields a 128-dimensional activation vector **z** with elements *z_i_* corresponding to the activation values of each hidden unit within the fully connected layer (second-to-last layer) that are recorded as the input image is passed through the network. We subsequently project all activation vectors onto an orthogonal basis computed via principal component analysis (PCA) that takes exclusively into consideration the activation vectors of (non-augmented) images used during training. Principal components explaining 99% of the variance define the new feature sub-space onto which all activation vectors are typically projected.

Feature vectors belonging to the fields-of-view (FOVs) of a well are aggregated by taking the element-wise median of the vectors. The resulting vector is taken as the feature vector representing the corresponding well. Likewise, to construct the feature vector for a given treatment, feature vectors of the treatment’s replicate wells are summarized by taking the element-wise median of the vectors.

### Distance and similarity calculations

To compare treatments, we use the cosine distance between two feature vectors. The cosine distance is defined as one minus the cosine of the angle between the vectors. The values thus range from 0 (denoting an identical direction for both vectors) to 2 (denoting opposite directions). To obtain a measure of *similarity* between treatments within the same numerical range, we subtract each cosine distance value from two.

### Clustering

We compute cosine distances among all pairs of treatments in a dataset. We use a hierarchical clustering algorithm to group treatments based on these pairwise cosine distance values. The resulting hierarchical tree is partitioned with a threshold value equivalent to the cosine distance entailed by an angle of *π*/3.

### Nearest neighbor classifier

Using pairwise cosine distances, we build a nearest neighbor classifier to investigate whether the feature vectors obtained via the unsupervised model encoded information that supported the retrieval of known phenotypic categories that had been manually assigned to a subset of treatments. Evaluation of the classifier’s performance requires splitting the feature vectors onto a training set and a test set. Given a feature vector from the test set, we determine its closest feature vector (i.e., its nearest-neighbor) within the training set, and assign the nearest neighbor’s phenotypic or MoA category to the test feature vector.

In the KiMorph dataset, we use a random hold-out cross-validation strategy where we randomly group all feature vectors into a training set and test set. The proportion of treatments assigned to the training set is 90%. Using the nearest-neighbor classifier, we predict the phenotype of the feature vectors in the test set, and evaluate the classification performance. We repeat the partitioning and evaluation process 50 times. The confusion matrix aggregating the results over the 50 repeats is shown in **Supplementary Table 1**.

In the BBBC021 dataset, we use a leave-one-compound-out validation strategy, where the training dataset excludes feature vectors of treatments (i.e., compound-concentration pairs) sharing the same compound as the test feature vector. We use all 103 treatments as test feature vectors once, obtain a nearest-neighbor prediction for the MoA, and compare the prediction with treatments’ known MoA. The resulting confusion matrix is shown in **Supplementary Table 4**.

### Image pre-processing

All image intensities are subjected to an Anscombe transform. Histogram normalization of each image is carried out on per-plate basis as described previously(16). All image intensities are mapped to an 8-bit range.

### Image datasets

The KiMorph dataset is available from the Wolfgang Huber Group EBI website at https://www.ebi.ac.uk/huber-srv/cellmorph/kimorph/.

The BBBC021 version 1 image dataset is available from the Broad Bioimage Benchmark Collection at http://www.broadinstitute.org/bbbc/BBBC021/.

Detailed description of the datasets can be found on their corresponding webpages.

## Acknowledgements

We would like to thank Florian Fuchs for fruitful discussions as well as for providing the KiMorph image data via the Wolfgang Huber Group EBI website. We also acknowledge the BBBC021 dataset provided by Peter D. Caie via the Broad Bioimage Benchmark Collection.

## Author Contributions

All authors conceived jointly the study. WJG designed the neural network architecture as well as training scheme, and performed the analysis. IH set up the deep learning computational framework. WJG and XZ wrote the manuscript.

## Supplementary Material

**Supplementary Table 1:** Confusion matrix on control siRNA treatments of the KiMorph dataset.

**Supplementary Table 2:** Distance matrix of entire siRNA collection of the KiMorph dataset with entries sorted according to an optimal leaf ordering for the hierarchical cluster tree computed based on the distance values

**Supplementary Table 3:** Clustering results of entire kinase siRNA collection of the KiMorph dataset

**Supplementary Table 4:** GO term enrichment analysis on the entire kinase siRNA collection of the KiMorph dataset

**Supplementary Table 5:** Confusion matrix on annotated treatments of the BBBC021 dataset

**Supplementary Table 6:** Similarity values of all treatments to reference treatments of the BBBC021 dataset

**Supplementary Table 7:** Distance matrix of entire compound collection of the BBBC021 dataset with entries sorted according to an optimal leaf ordering for the hierarchical cluster tree computed based on the distance values

**Supplementary Table 8:** M-CNN architecture used to analyze both the KiMorph and BBBC021 dataset

**Supplementary Figure 1:** Similarity-vs-concentration curves for selected compounds of the BBBC021 dataset.

**Supplementary Figure 2:** Example images of Cochicine at varying concentrations.

**Supplementary Software**: Solver definition and network specification files.

